# Evasion of neutralizing antibodies by Omicron sublineage BA.2.75

**DOI:** 10.1101/2022.07.19.500716

**Authors:** Daniel J. Sheward, Changil Kim, Julian Fischbach, Sandra Muschiol, Roy A. Ehling, Niklas K. Björkström, Gunilla B. Karlsson Hedestam, Sai T. Reddy, Jan Albert, Thomas P. Peacock, Ben Murrell

## Abstract

An emerging SARS-CoV-2 Omicron sublineage, BA.2.75, is increasing in frequency in India and has been detected in at least 15 countries as of 19 July 2022. Relative to BA.2, BA.2.75 carries nine additional mutations in spike. Here we report the sensitivity of the BA.2.75 spike to neutralization by a panel of clinically-relevant and pre-clinical monoclonal antibodies, as well as by serum from blood donated in Stockholm, Sweden, before and after the BA.1/BA.2 infection wave.

BA.2.75 largely maintains sensitivity to bebtelovimab, despite a slight reduction in potency, and exhibits moderate susceptibility to tixagevimab and cilgavimab. For sera sampled both before and after the BA.1/BA.2 infection wave, BA.2.75 does not show significantly greater antibody evasion than the currently-dominating BA.5.

## Main Text

Towards the end of 2021, SARS-CoV-2 vaccine effectiveness was threatened by the emergence of the Omicron clade (B.1.1.529), with more than 30 mutations in spike. Recently, several sublineages of Omicron have demonstrated even greater immune evasion^1–4^, and are driving waves of infections across the globe.

One emerging sublineage, BA.2.75, is increasing in frequency in India and has been detected in at least 15 countries as of 19 July 2022. Relative to BA.2, BA.2.75 carries nine additional mutations in spike (Fig. S1): K147E, W152R, F157L, I210V, G257S, G339H, G446S, N460K, and a reversion towards the ancestral variant, R493Q. G446S has been predicted to be a site of potential escape from antibodies elicited by current vaccines that still neutralize Omicron^5^. Further, it has been identified as a site of potential escape from LY-CoV1404 (bebtelovimab), which represents one of the last remaining classes of first-generation monoclonal antibodies that are still able to cross-neutralize BA.2 and BA.4/5^1^. As waves of Omicron infections have occurred in many countries, identifying the sensitivity of newly emerging variants to neutralization by sera sampled subsequent to these waves is required to inform public health policy.

Here we report the sensitivity of the BA.2.75 spike to neutralization by a panel of clinically-relevant and pre-clinical monoclonal antibodies, as well as by serum from blood donated in Stockholm, Sweden during week 45, 2021, (N=20) and week 15, 2022, (N=20) This coincides with points before and after a large wave of infections dominated by BA.1 and BA.2 (Dec 2021 - Feb 2022), as well as an expansion of vaccine ‘booster’ doses (Fig. S2).

Cilgavimab had approximately 11-fold reduced potency against BA.2.75 compared to the ancestral B.1 (D614G), in line with its potency against BA.5 (Fig 1A and Fig S3). While only capable of extremely weak neutralization of BA.2, tixagevimab saw partially restored activity against BA.2.75, possibly due, in part, to the reversion to the ancestral amino acid at spike position 493. While bebtelovimab indeed demonstrated reduced potency against BA.2.75, likely due to G446S, the loss was only around 7-fold, and bebtelovimab still potently neutralizes BA.2.75. Casivirimab, imdevimab, bamlanivimab, and etesevimab all failed to neutralize BA.2.75. These titers are largely concordant with recent data from others^6,7^, though the magnitude of the loss of potency for cilgavimab/COV2-2130 shows substantial variation between the three studies.

**Figure 1.**
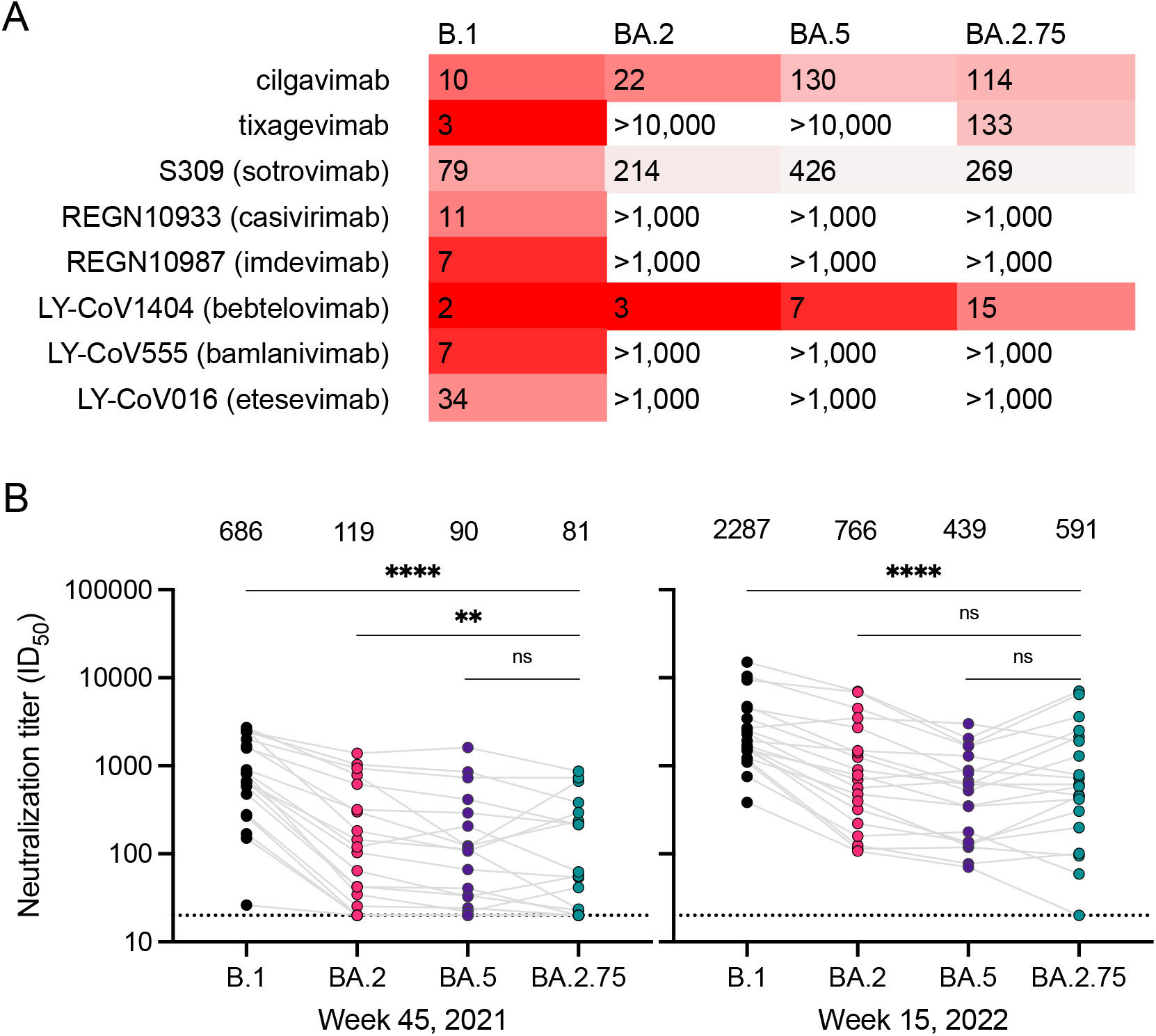
**A.** Neutralizing IC50 (50% inhibitory concentration) titers (ng/μl) for monoclonal antibodies against ancestral B.1 (D614G) and Omicron sublineages BA.2, BA.5 and BA.2.75 in a pseudovirus neutralization assay. **B.** Neutralization of BA.2.75 relative to BA.2, BA.5 and B.1 by serum (N=20) from blood donated in week 45, 2021 (8 Nov - 14 Nov) in Stockholm, Sweden, prior to a wave of infections dominated by BA.1 and BA.2 (**left**). Neutralization by serum (N=20) donated in week 15, 2022 (11 Apr - 17 Apr), after the infection wave (**right**). Depicted above are the geometric mean ID50 (50% inhibitory dilution) titres. Serum with an ID50 less than the lowest dilution tested (20, dotted line) is plotted as 20.

BA.2.75 was neutralized with the lowest geometric mean titer of all variants evaluated by ‘prewave’ sera (Fig. 1B), with titers to BA.2.75 approximately 8-fold reduced compared to ancestral B.1 (D614G). Pre-wave, titers against BA.2.75 were slightly but significantly lower than those against BA.2, and comparable to those against BA.5. Sera sampled following the BA.1/BA.2 infection wave displayed substantially improved neutralization against ancestral B.1 as well as enhanced cross-neutralization of omicron sublineages. Geometric mean titers against BA.2.75 were more than 7-fold higher for ‘post-wave’ compared to ‘pre-wave’ sera (Fig. S4), likely reflecting a combined contribution of BA.1 and BA.2 infections, as well as 3rd dose booster vaccine rollout, with coverage in Stockholm expanding among persons 18 years or older from 5.1% in week 45 2021 to 59% in week 15 2022 (Fig. S2). The relative sensitivity of BA.2.75 in these cohorts of blood donors is largely concordant with those seen in recipients of CoronaVac with and without BA.1/BA.2 breakthrough infection^7^.

As infection histories become more complex, and a large proportion of infections go undetected, monitoring of population-level immunity from random samples is increasingly critical for understanding and contextualizing the immune evasion properties of new variants. Here we show that the currently dominating BA.5, and the emerging sublineage BA.2.75 are similarly resistant to neutralization by serum from randomly sampled seropositive blood donations from Stockholm. BA.2.75 largely maintains sensitivity to bebtelovimab despite a slight reduction in potency, and exhibits moderate susceptibility to tixagevimab and cilgavimab. The sensitivity of BA.2.75 to neutralization by antibody classes that BA.5 has escaped suggests there is significant scope for further escape in lineages of BA.2.75.

## Appendix

### Supplementary Figures and Tables

**Figure S1.**
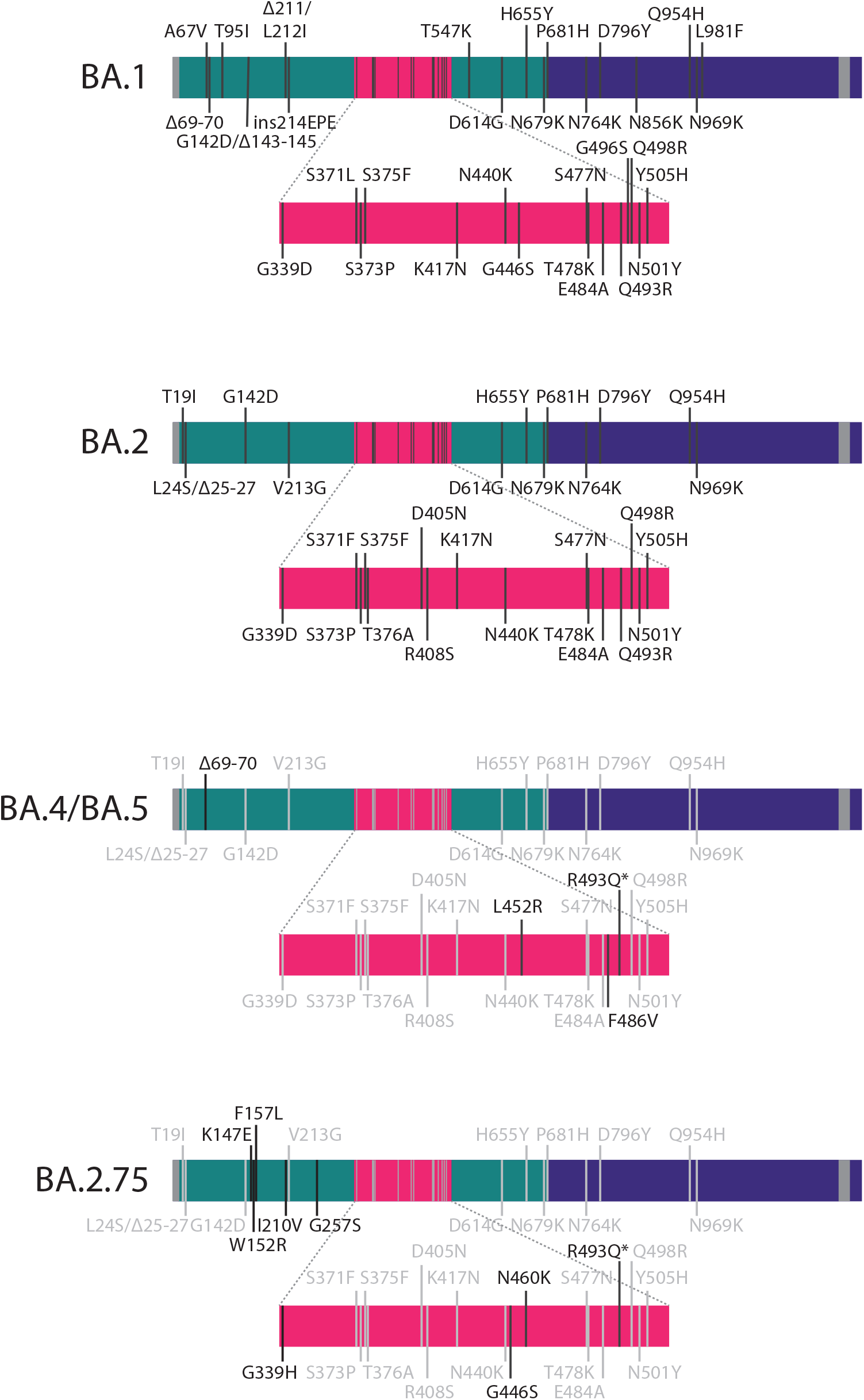
List of spike mutations in Omicron sub-variants BA.1, BA.2, BA.4/5, and BA.2.75. *indicates a reversion to the ancestral amino acid.

**Figure S2.**
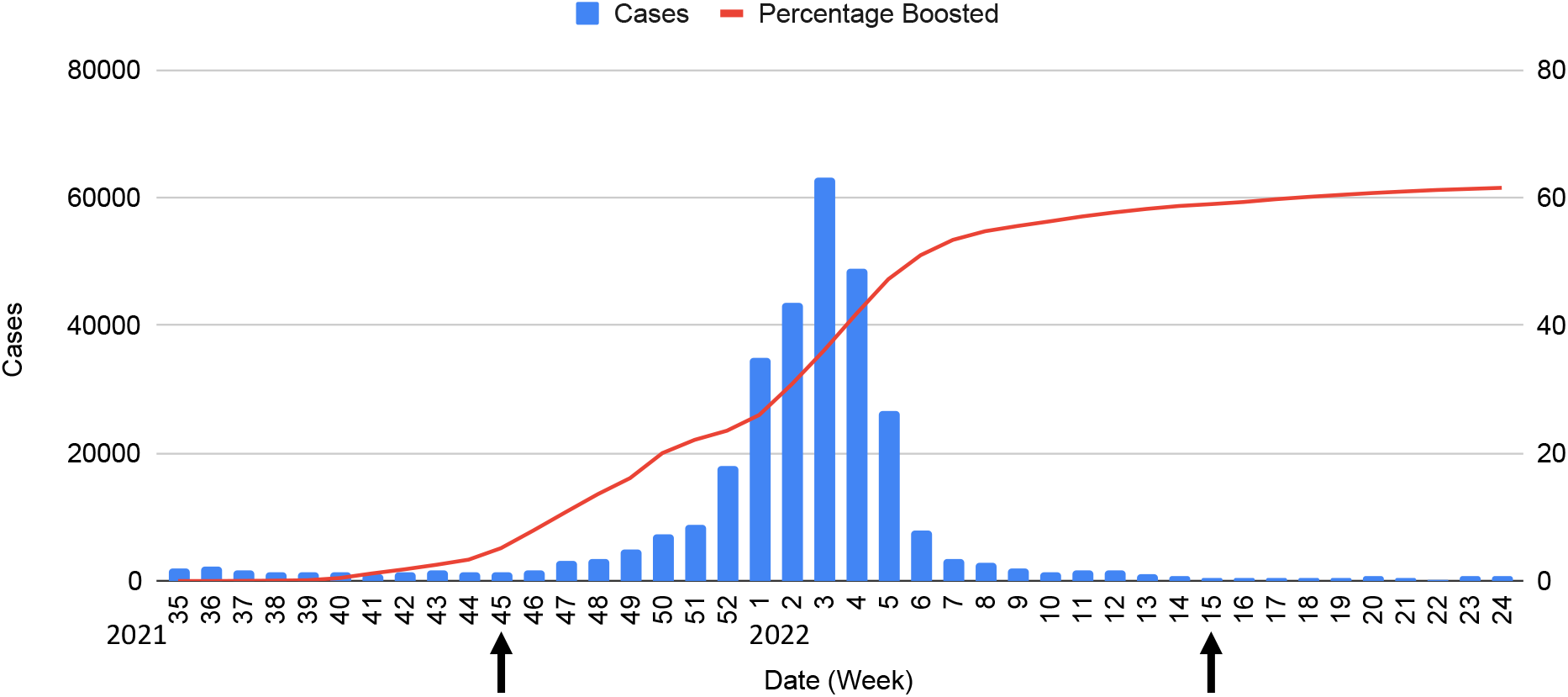
Timecourse of SARS-CoV-2 infections and vaccinations in Stockholm, Sweden. Depicted are numbers of confirmed SARS-CoV-2 cases in Stockholm per week^1^ (blue), as well as cumulative percentage of individuals, 18 and older, who had received their 3rd dose “booster” vaccination^2^ (red). The date range spans Sweden’s first Omicron infection wave, which was initially dominated by BA.1, but followed by BA.2 cases^3^. Black arrows denote the weeks from which our serum samples were obtained. The first sampling point is prior to any confirmed Omicron infections, and the second is after the bulk of BA.1 and BA.2 infections, but prior to the arrival of BA.4 or BA.5.

**Figure S3.**
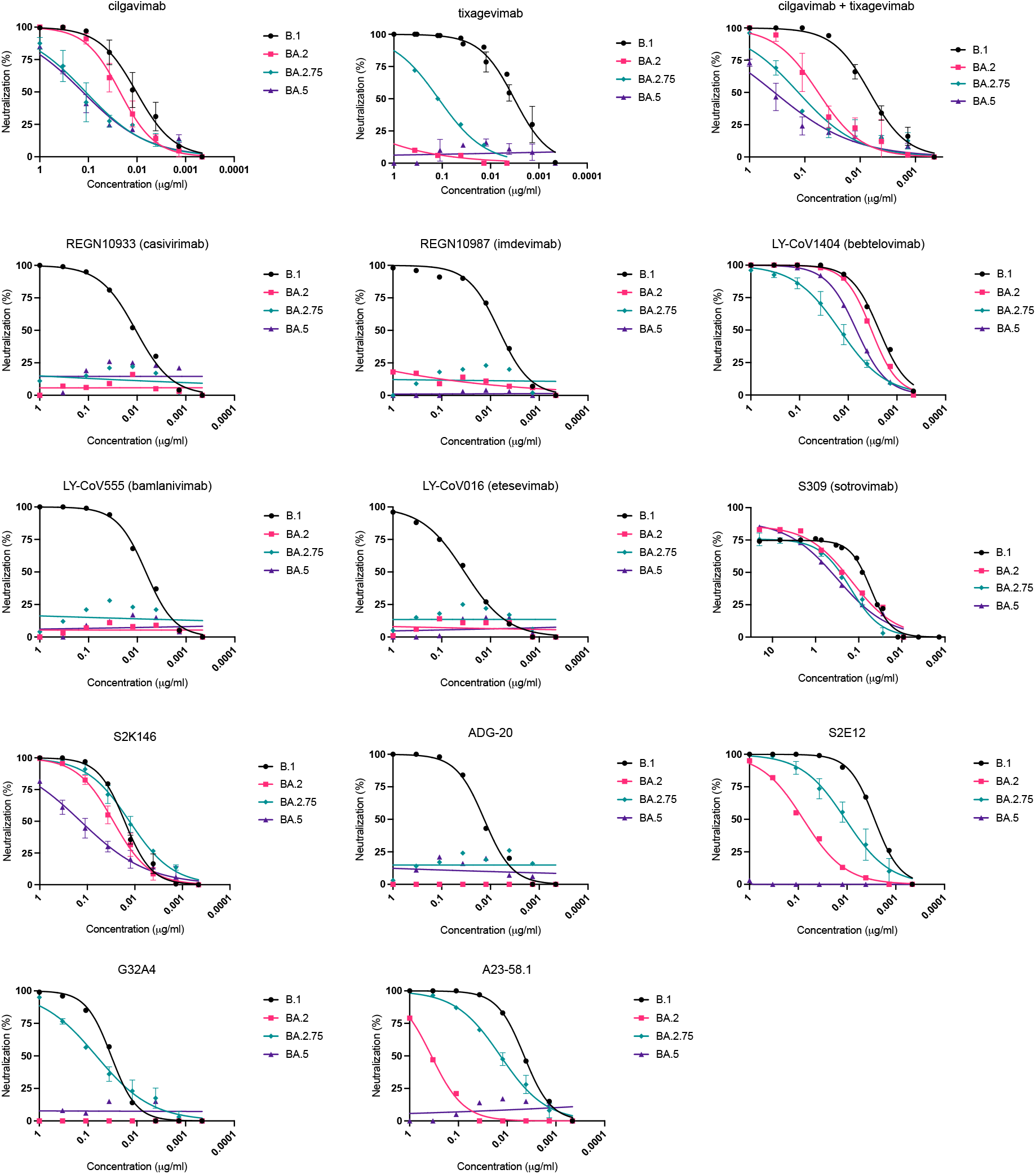
Neutralization curves for monoclonal antibodies against B.1 (D614G), and Omicron sublineages BA.2, BA.5 and BA.2.75.

**Table S1.** Relative sensitivity of BA.2.75 to therapeutic and pre-clinical monoclonal antibodies. IC_50_ titers (50% inhibitory concentration, in ng/μl) in a pseudovirus neutralization assay are tabulated for monoclonal antibodies against B.1 and Omicron sublineages BA.2, BA.5 and BA.2.75.

|  | B.1 | BA.2 | BA.5 | BA.2.75 |
| --- | --- | --- | --- | --- |
| cilgavimab<sup>4</sup> | 10 | 22 | 130 | 114 |
| tixagevimab<sup>4</sup> | 3 | >1,000 | >1,000 | 133 |
| S309 (sotrovimab)<sup>5</sup> | 79 | 214 | 426 | 269 |
| REGN10933 (casirivimab)<sup>6</sup> | 11 | >1,000 | >1,000 | >1,000 |
| REGN10987 (imdevimab)<sup>6</sup> | 7 | >1,000 | >1,000 | >1,000 |
| LY-CoV1404 (bebtelovimab)<sup>7</sup> | 2 | 3 | 7 | 15 |
| LY-CoV555 (bamlanivimab)<sup>8</sup> | 7 | >1,000 | >1,000 | >1,000 |
| LY-CoV016 (etesevimab)<sup>9</sup> | 34 | >1,000 | >1,000 | >1,000 |
| S2E12<sup>10</sup> | 3 | 79 | >1,000 | 8 |
| S2K146<sup>11</sup> | 15 | 35 | 180 | 15 |
| ADG20<sup>12</sup> | 13 | >1,000 | >1,000 | >1,000 |
| A23-58.1<sup>13</sup> | 4 | 358 | >1,000 | 13 |
| G32A4<sup>14</sup> | 34 | >1,000 | >1,000 | 78 |

**Fig S4.**
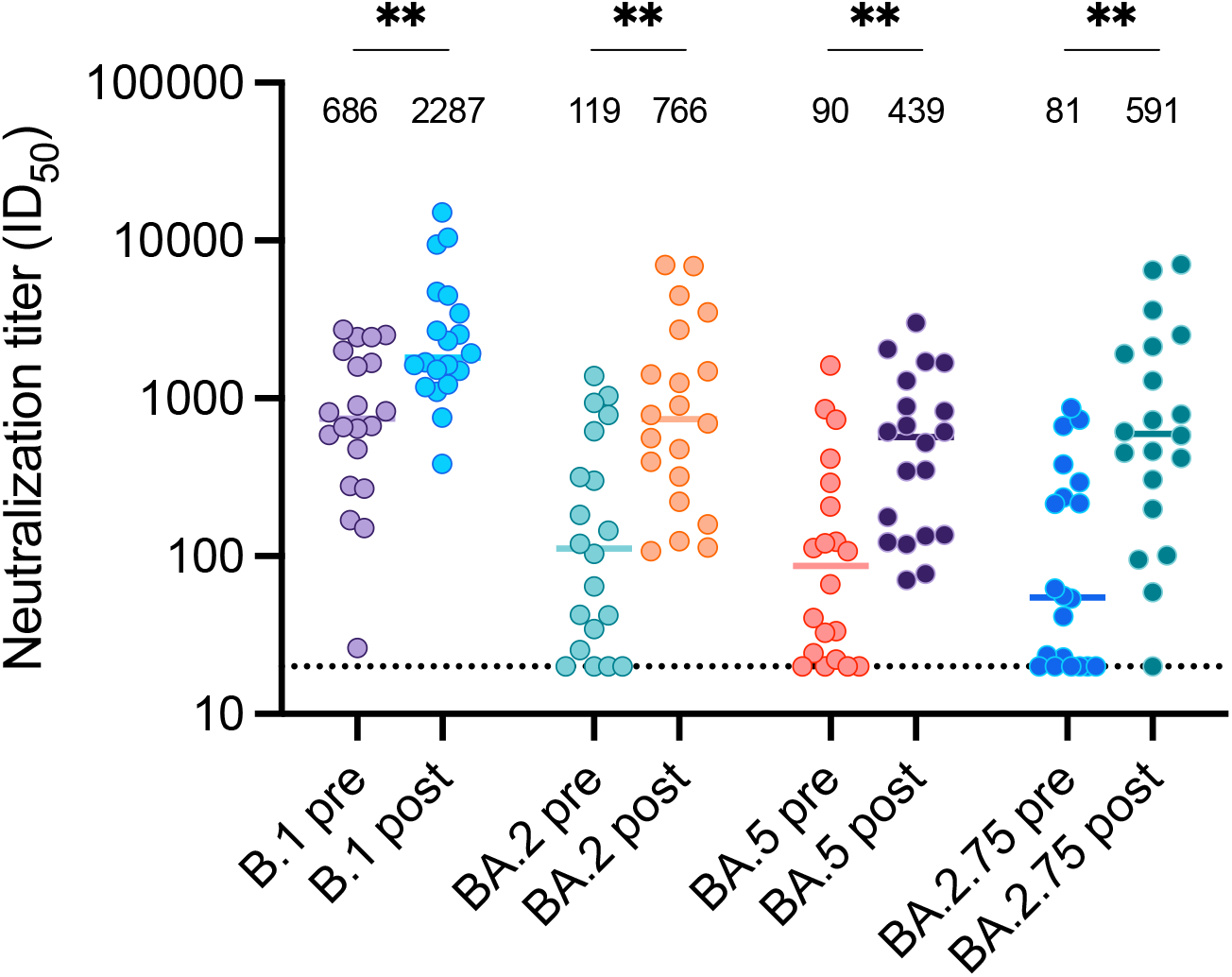
Comparison of ‘pre-wave’ and ‘post-wave’ neutralizing titers. Neutralizing ID50s (50% inhibitory dilution) are shown for serum from blood donated in Stockholm, Sweden during week 45, 2021 (8 Nov - 14 Nov), prior to a wave of infections dominated by BA.1 and BA.2 (**pre**), and week 15, 2022 (11 Apr - 17 Apr), after the infection wave (**post**). Depicted above are the geometric mean ID50 titres. Sera with ID50 less than the lowest dilution tested (20, dotted line) are plotted as 20. ** P<0.01; *** P<0.001 (Mann-Whitney U test, corrected for multiple comparisons using the Holm procedure).

### Methods

#### Cell culture

HEK293T cells (ATCC CRL-3216) and HEK293T-ACE2 cells (stably expressing human ACE2) were cultured in Dulbecco’s Modified Eagle Medium (high glucose, with sodium pyruvate) supplemented with 10% fetal bovine serum, 100 units/ml Penicillin, and 100 μg/ml Streptomycin. Cultures were maintained in a humidified 37°C incubator (5% CO_2_).

#### Pseudovirus Neutralization Assay

Pseudovirus neutralization assay was performed as previously^15^. Briefly, spike-pseudotyped lentivirus particles were generated by co-transfection of HEK293T cells with a relevant spike plasmid, an HIV gag-pol packaging plasmid (Addgene #8455), and a lentiviral transfer plasmid encoding firefly luciferase (Addgene #170674) using polyethylenimine. The BA.2.75 spike plasmid was generated by introducing the following mutations into the BA.2 spike, by multisite directed mutagenesis: K147E, W152R, F157L, I210V, G257S G339H, G446S, N460K, R493Q, which was subsequently confirmed by sequencing.

Neutralization was assessed in HEK293T-ACE2 cells. Pseudoviruses sufficient to produce ±100,000 RLU were incubated with serial 3-fold dilutions of serum for 60 minutes at 37°C in a black-walled 96-well plate. 10,000 HEK293T-ACE2 cells were then added to each well, and plates were incubated at 37°C for 48 hours. Luminescence was measured using Bright-Glo (Promega) on a GloMax Navigator Luminometer (Promega). Neutralization was calculated relative to the average of 8 control wells infected in the absence of serum.

#### Monoclonal antibodies

Cilgavimab and tixagevimab were evaluated as their clinical formulations. For the rest of the monoclonal antibodies evaluated, antibody sequences were extracted from deposited RCSB entries, synthesized as gene fragments, cloned into pTWIST transient expression vectors by Gibson assembly or restriction cloning, expressed and purified, all as previously described^16^.

#### Serum samples

Serum samples from anonymized blood donors from Stockholm, Sweden, were obtained from week 45, 2021 (prior to the BA.1/BA.2 Omicron infection wave), and from week 15, 2022 (after the BA.1/BA.2 Omicron infection wave, but prior to the arrival of BA.4 or BA.5); see Fig S2. 25 serum samples from each time point were pre-screened for detectable neutralization activity against ancestral B.1 (D614G), and 20 samples with detectable activity against B.1 (D614G) for each time point were selected, randomly, for this study. Sera were heat inactivated at 56°C for 60 minutes prior to use in neutralization assays.

#### Ethical Statement

The blood donor samples were anonymized, and not subject to ethical approvals, as per advisory statement 2020–01807 from the Swedish Ethical Review Authority.

#### Statistical analysis

Individual ID50 and IC50 values for each sample against each variant were calculated in Prism v9 (GraphPad Software) by fitting a four-parameter logistic curve to neutralization by serial 3-fold dilutions of serum/antibody. Comparison of titers between variants was assessed using paired Wicoxon signed-rank tests. Comparison of titers pre- and post-wave was assessed using unpaired Mann-Whitney tests. Correction for multiple comparisons was performed using the Holm procedure^17^, implemented in MultipleTesting.jl, in the Julia language for Scientific Computing. P values are summarized as: ns P>0.05; * P<0.05; ** P<0.01; *** P<0.001; **** P<0.0001.

## Author contributions

Conceptualization, D.J.S., T.P.P., B.M.;

Formal analysis, D.J.S., B.M.;

Conducted the assays, D.J.S., C.K., J.F., T.P.P.;

Designed the methodology, D.J.S., C.K., R.E., T.P.P, B.M.;

Responsible for figures and tables, D.J.S., T.P.P., B.M.;

Resources, S.M., R.E., S.R, N.K.B., G.B.K.H., J.A., B.M.;

Oversaw the study, D.J.S., G.B.K.H., S.T.R., J.A., B.M;

Funding Acquisition, S.T.R., G.B.K.H., J.A., T.P.P., B.M.

Writing – original draft, D.J.S.;

Writing – review & editing, D.J.S., J.F., G.B.K.H., J.A., T.P.P., B.M.

D.J.S and B.M. were responsible for the decision to submit the manuscript for publication.

## Acknowledgements

pCMV-dR8.2 dvpr was a gift from Bob Weinberg (Addgene plasmid # 8455; http://n2t.net/addgene:8455; RRID:Addgene_8455). pBOBI-FLuc was a gift from David Nemazee (Addgene plasmid # 170674; http://n2t.net/addgene:170674; RRID:Addgene_170674). We also wish to thank the researchers, globally, who contributed to the sequencing and identification of the emerging BA.2.75 variant.

